# Enrichment of polyphenol metabolism within the ubiquitous SAR116 ‘Puniceispirillales’ bacterioplankton

**DOI:** 10.64898/2026.07.17.739263

**Authors:** Jordan T. Coelho, Mikayla A. Borton, J. Cameron Thrash

## Abstract

Marine dissolved organic matter (DOM) is a complex pool of substrates, and a central challenge in marine biogeochemistry is to determine which DOM compounds are produced, consumed, or exchanged by microorganisms. Aromatic compounds, including polyphenols, comprise an important chemical class of marine DOM generated both *in situ* and also supplied via atmospheric deposition, terrestrial runoff, or natural seeps. Here we demonstrate, through pangenomics and metabolic reconstruction, the enrichment of genes for degradation of a wide variety of polyphenols in the cosmopolitan SAR116 bacterioplankton (Puniceispirillales). Gene and pathway conservation, the evolutionary history of key ring-cleaving dioxygenases, and expression patterns from global ocean surveys support the conclusion that these transformations represent core elements of SAR116 ecophysiology. The degradation pathway for protocatechuate, an end member for many polyphenol transformations, occurred in all subclades, and we found evidence of different evolutionary histories for genes in this pathway, including gene loss, analogous substitution, and retention of horizontally transferred genes from the *Gammaproteobacteria.* Furthermore, variation in these and other pathway-specific gene losses suggest that some SAR116 polyphenol metabolism may occur through a division of labor in concert with other community members and/or require novel or promiscuous enzymes. The enrichment and expression of polyphenol degradation pathways define a previously underappreciated metabolic niche for SAR116 and provides new evidence emphasizing the importance of polyphenol metabolism in the marine carbon cycle.

## Introduction

Marine dissolved organic matter (DOM) comprises a complex mixture of molecules that are produced, consumed, and altered by microbes [1–3], and also sustain the microbial loop [4, 5]. The bulk of marine DOM is produced by phytoplankton [3, 6], and DOM consumption by heterotrophic microbes can often provide insight into ecosystem processes and biogeochemical cycles [1, 7–13]. The DOM pool consists of thousands to tens of thousands of distinct organic substrates [14, 15], and despite incredible advances in analytical chemistry, we still face many challenges in characterizing this vastly complex collection of compounds [16].

Microbes can help us determine DOM components of interest based on the potential for, or realized measurements of, their ability to metabolize, uptake, exchange, or generate certain compounds. For example, exometabolomics of multiple important phytoplankton has highlighted a suite of secreted compounds important for future study, including some that are underappreciated in phycospheres such as glutamic acid, chitobiose, and 2’deoxyuridine [17]. Comparative genomics and experimental physiology of SAR11 revealed the importance of one-carbon and methylated compounds for energy metabolism in this group [18]. *In situ* measurements in the Gulf of Mexico demonstrated that *Prochlorococcus* and *Synechococcus* spp. produced hydrocarbons, specifically n-pentadecane, that sustained growth of an enriched community of known hydrocarbon degraders and revealed a novel metabolic niche for Marine Group II (MGII) Archaea [19]. These few examples highlight how microbiology helps inform chemical oceanography, specifically in the search for critical components of the DOM pool.

We were drawn to a collection of observations about DOM metabolism that intersected with our ongoing investigations into the ecology of the ubiquitous SAR116 bacterioplankton. SAR116 Alphaproteobacteria (Puniceispirillales) are characteristic surface ocean microorganisms, typically representing about 5-12% of the prokaryotic community [20–25]. Obligately aerobic, they also perform heterotrophic [26–28] and a few lithotrophic [28, 29] metabolisms, in particular degrading amino acids [30], five-carbon carbohydrates [30], DMSP [31], and other low molecular weight DOM [26, 28]. They frequently associate with phytoplankton blooms [32–34] and especially late-bloom phases [33]. SAR116 have also been enriched in seawater incubations with added naphthalene from the Southern California Bight [35], and metatranscriptomic analysis from the Sapelo Island Microbial Observatory on the Georgia, USA coast highlighted benzoate degradation to be an indicator gene for SAR116 [30]. These observations suggested the role of SAR116 members in degradation of aromatic compounds. However, SAR116 are not typically observed as primary or successional responders in oil spills, or inhabitants of natural oil seeps [36–41].

Motivated by these seemingly incongruous observations about their relationship with aromatic compounds, we investigated the potential for SAR116 to degrade them, and here present evidence that members of SAR116 consistently encode pathways for metabolism of polyphenols. Polyphenols are classically defined as plant-derived secondary metabolites [42], however there is evidence that they are autochthonous to the marine systems [43–54]. The roles of polyphenols in microbial communities have been evaluated in terrestrial systems such as soils [55–57], permafrost [42], and gut and rumen systems [58, 59], however the relevance of polyphenols to microbial metabolism in marine systems, particularly open ocean systems, remains largely unknown. Using comparative genomics, phylogenetics, and metatranscriptomic recruitment, we predict that polyphenolic substrates are important components of DOM used by the SAR116 clade. Given their global distribution as well as their evolutionary and habitat diversity, the emphasis and variation of polyphenol metabolism within SAR116 helps us better understand their niche in the global oceans and highlights the importance of polyphenols across different marine ecosystems.

## Materials and Methods

### Pangenomics and identification of aromatic carbon genes

We analyzed a pangenome comprising 349 genomes [28] (Table S1) including recently isolated [21, 60] and sequenced [28] SAR116 isolate genomes and publicly available SAR116 genomes [61–64]. Genomes were filtered by quality metrics, dereplicated, and assigned to the same phylogenomic groups as previously described [28, 65, 66]. In this work, using the prior pangenomic analysis [28], we manually examined the SAR116 pangenome summary (Table S2) for aromatic and polyphenolic degradation genes via Pfam, KEGG Orthology, TIGRFAM, Prosite, and PANTHER accession numbers (Tables S3, S4). We also updated our prior annotations with DRAM v1.5.0 [67], including the CAMPER (Curated Annotations for Microbial Polyphenol Enzymes and Reactions) module [42, 68]. Annotations were then distilled in DRAM to generate the metabolism_summary.xls (Table S6). We used the CAMPER distillate output to estimate pathway completion per genome for 100 polyphenol related pathways, accounting for the number of subunits and number of pathway steps. For Figure 1, genomes were considered to have a pathway if it was ≥ 50% complete, and we cross-referenced the CAMPER annotations (Table S5, S6) with the pangenomic annotations from our previous study (Table S1). For specific pathways (gentisic acid, homogentisic acid, and protocatechuic acid degradation), we only considered the pathway complete if the ring-cleaving dioxygenase was detected.

**Figure 1:**
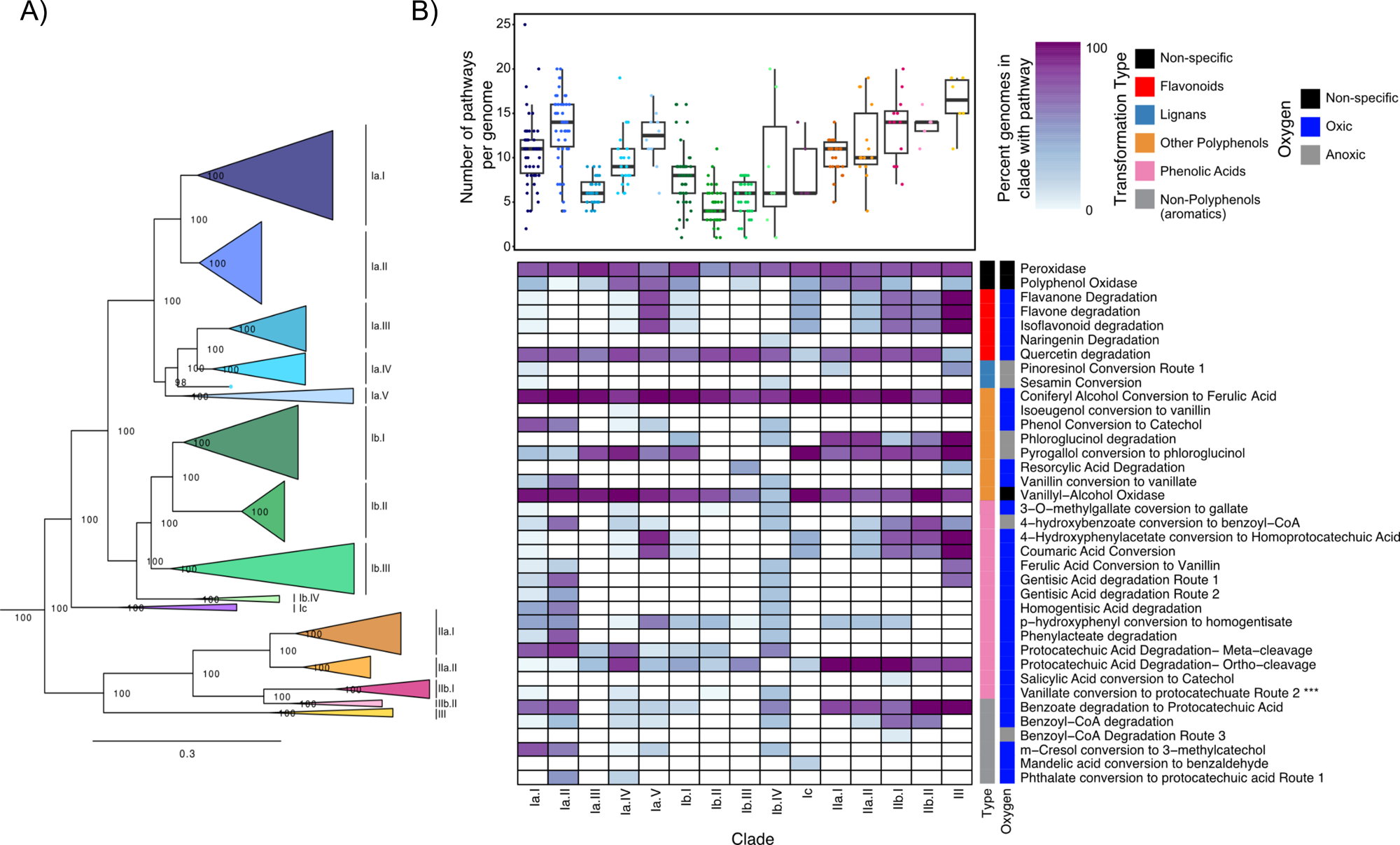
CAMPER Polyphenol transformation across SAR116 genomes. (A) Phylogenomic tree from our previous study, delineating SAR116 subclades [28] (B) CAMPER output by subclades from A. The boxplot (top) shows the number of polyphenol transformation/degradation pathways per genome, grouped by subclade, with each datapoint representing a genome. Boxplots represent the median and one standard deviation. The heatmap (bottom) shows the percentage of genomes containing a pathway, by pathway type. Rows represent pathways, and columns represent subclades. Darker colors indicate higher percent representation of a pathway within a subclade. Pathways are defined by both the type of transformation based on polyphenol category, and the oxygen requirement for the pathway. Vanillate conversion to protocatechuate (Pfam, PF19112, Tables S2, S4) is labeled with three asterisks (***).

### Single Gene Phylogenetics

We performed phylogenetic inference of aromatic ring-cleaving dioxygenase amino acid sequences found in SAR116 genomes to understand their evolutionary history and better validate the predicted function of their corresponding genes. Gentisate 1,2-dioxygenase (SdgD), catechol 2,3-dioxygenase (DmpB), hydroxyquinol 1,2-dioxygenase (ChqB), and the active site-encoding subunits for protocatechuate 4,5-dioxygenase (LigB), and protocatechuate 3,4-dioxygenase (PcaH) [69] were used in the analysis of aromatic ring-cleaving dioxygenase genes. Amino acid sequences of the transcriptional regulator, LysR, and downstream protocatechuate degrading enzyme, LigJ, were also included for phylogenetic inference to evaluate horizontal gene transfer. Separately for each ring-cleaving dioxygenase gene, we searched all SAR116 amino acid sequences against the NCBI RefSeq database [70] via BLAST [71]. For each SAR116 query sequence, we selected the top 25 best hits and then filtered all results to remove duplicate sequences. Including the SAR116 query sequences, this resulted in 157 SdgD, 527 DmpB, 548 ChqB, 275 LigB, and 351 PcaH amino acid sequences for phylogenetic inference. We separately aligned fasta files for all homologous protein groups with MUSCLE [72] and trimmed with TrimAl [73], both with default settings. We performed maximum-likelihood inference with IQ-TREE v2.1.2 [74] using the automated amino acid substitution best-fit model estimator ‘-m MFP’ and traditional bootstrapping with 1000 replicates. The resulting trees were visualized with custom R-scripts (available on FigShare) using *Ggtree* v3.2.1 and *Treeio* v1.18.1 R packages [75–77], rooted at the midpoint, and nodes were ordered in increasing order.

### Protocatechuate ring-cleaving dioxygenase gene neighborhood and synteny

We analyzed the gene neighborhoods and synteny of the protocatechuate ring-cleaving dioxygenase genes to further investigate their evolutionary history. Using the Anvi’o v7.1 [78] generated “gene_callers_id” from the pangenome annotation summary (Table S2), gene annotations for each genome were ordered chronologically using a custom R-script (available on FigShare) to reflect a syntenic sequence. We examined the neighborhoods 10 genes up- and downstream from the *pcaGH* and *ligAB* dioxygenase genes. We used the ‘anvi-export-gene-calls’ function in Anvi’o v7.1 to determine gene directionality, whether the recorded upstream and downstream genes were on the same contig as the dioxygenase genes, the size of the genes, and the intergenic spacer lengths. Only contigs where upstream and downstream genes formed a contiguous sequence with the dioxygenase genes were retained (Table S7). These contiguous sequences were compared across subclades. With the information above, a representative sequence from a genome with the highest completion from each subclade was selected for further investigation of gene neighborhoods surrounding the dioxygenase genes.

### Biogeography of ring-cleavage gene expression

To qualitatively validate the degradation of polyphenolic and aromatic compounds by SAR116 members *in situ*, we competitively recruited metatranscriptomic reads from the planktonic size fraction (0.8*µ*m - 5*µ*m) of the TARA Oceans metatranscriptomic dataset [79] against all 349 SAR116 genomes. We used bowtie2 v2.5.2 for read mapping [80], SAMtools v.1.19.2 for read alignment processing [81], and coverM v0.7.0 [82] to filter and include only aligned reads that were greater than 95% identical and an alignment length of at least 75% of the read. Using ‘anvi-export-gene-calls’ we located the exact positions for ring-cleaving genes of interest and used bedtools v2.31.1 [83] to quantify reads that aligned to each loci (Table S8). The total coverage of aligned reads to genes of interest were summed according to the major subclade designation (I, II, or III) for each SAR116 genome, per metatranscriptomic sample.

### Data visualization

We visualized pangenomic, phylogenetic, gene synteny, and biogeography data using custom R-scripts (available on FigShare). For single gene phylogenetics, the resulting trees were visualized with *Ggtree* v3.2.1 and *Treeio* v1.18.1 R packages [75–77], rooted at the midpoint, and nodes were organized in increasing order. Gene synteny analyses were qualitatively visualized using *gggenomes* v1.0.1 [84]. Biogeographic distributions were processed and qualitatively visualized using *ggolot2* [85], *scatterpie* [86], and *maps* [87].

## Results and Discussion

### Polyphenol chemical transformations

We identified extensive potential for the transformation of polyphenols spanning the multiple subclades of SAR116 (Fig. 1A), with all genomes having at least one predicted polyphenol chemical transformation pathway, and an average of nine pathways per genome (Fig. 1B). Many of the enzymes in these pathways have specific oxygen requirements, and while most pathways encoded in SAR116 genomes require oxygen, some of these reactions can occur with or without oxygen, and others have only been documented in anoxic environments (Fig. 1B). There are two substrate-nonspecific genes typically described in polyphenol degradation [88–90], polyphenol oxidase and peroxidase (AA2, also involved in lignin degradation [91]), that both destabilize carbon linkages via the formation of free radicals [89, 92, 93]. These two genes were widespread across the SAR116 genomes (Fig. 1B), suggesting a general capacity for polyphenol metabolism among SAR116 members that is not substrate specific. Collectively, subclade III encoded the highest number of pathways for polyphenolic metabolism (Fig. 1B), although individual genomes in these clades had a wide range of pathway totals. For example, one member of the Ia.I clade, the cultured representative strain LSUCC0719, encoded 25 pathways for polyphenol transformations, exceeding the median of any one clade (average range across clades 4.2 −16.3). Conversely, other clades such as IIb.II and III had lower variability among members. The widespread distribution of polyphenolic degradation genes highlights polyphenols as likely core metabolites for SAR116.

The most common group of transformations encoded by SAR116 genomes was in the broad “Other Polyphenol” category, consisting of genes/pathways for interacting with small monomeric phenolic derivatives. For example, pathway representation for the transformation of coniferyl alcohol, a lignin precursor [94], to ferulate via CalAB (330/349 genomes), and vanillyl alcohol, a byproduct of lignin degradation [95], to vanillate via AA4 (293/349 genomes), were very common (Figs. 1B, 2). Additionally, subclades Ia.I, Ia.II, and Ib.IV encoded vanillin dehydrogenase (*vdh*) to convert vanillin to vanillate (Figs. 1B, 2). The potential for conversion of phenol to catechol via phenol 2-monooxygenase was widespread in subclades Ia.I and Ia.II, but was sparsely distributed in subclades Ia.IV, Ia.V, and Ib.IV (Figs. 1B, 2). Phenol 2-monooxygenase produces NADPH in the conversion of phenol to catechol [96], so SAR116 members may harvest the NAPDH from the reaction and then secrete catechol or alternatively linearize the catechol via a catechol 2,3-dioxygenase (*dmpB*, see below, Fig. 3).

**Figure 2:**
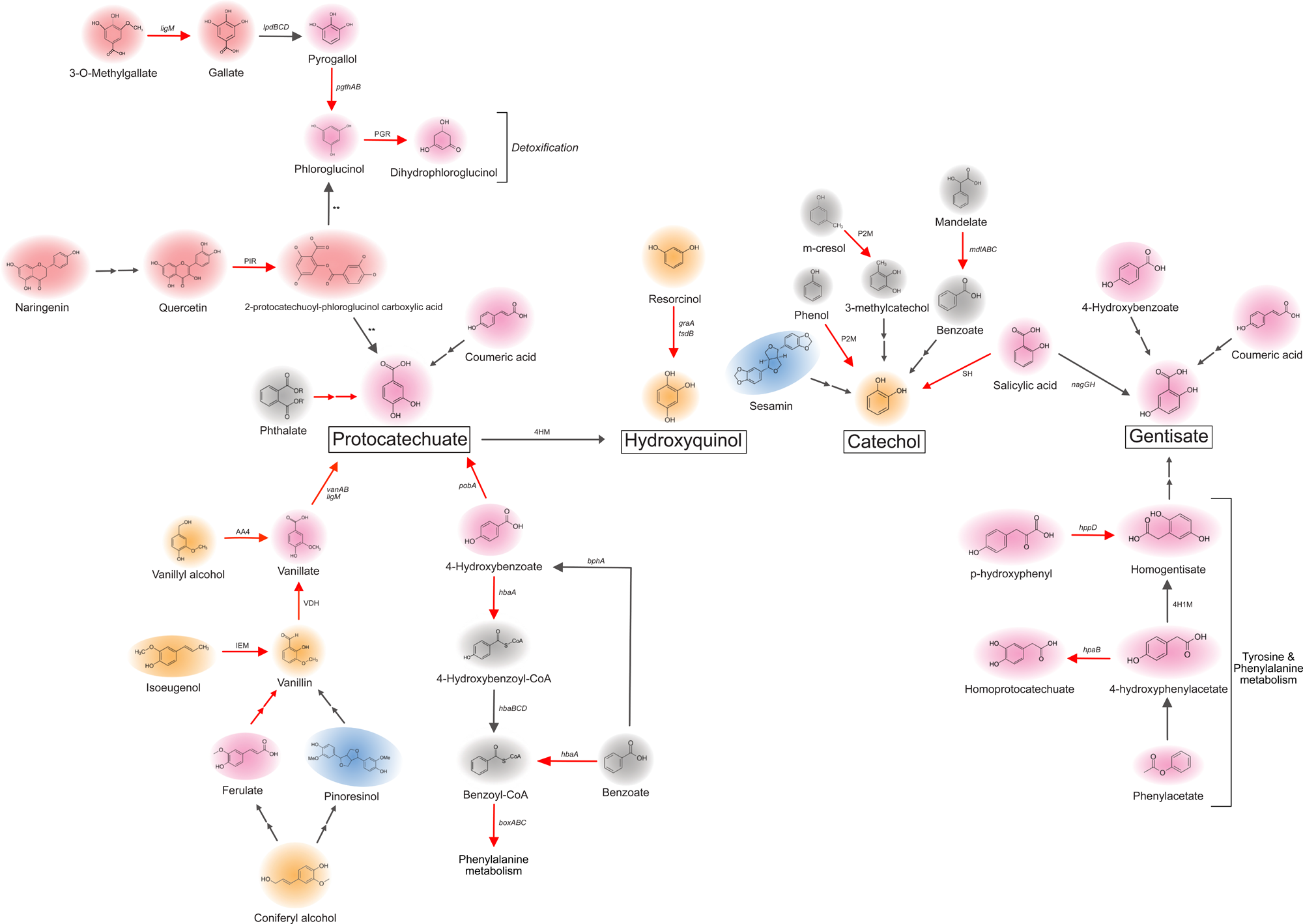
Conceptual schematic of polyphenolic funneling to terminal end-products. Chemical structures were selected based on the detection of transformation/degradation genes in SAR116 genomes, and are color coded based on their polyphenol category in Figure 1. Arrows indicate transformations that have been previously defined [120, 124–129], and represent the possible funneling of polyphenolic compounds to fuel protocatechuate, hydroxyquinol, catechol, and gentisate metabolism in SAR116 cells. Double arrows represent transformations that require multiple steps. ** = putative uncharacterized depsides/esterases that are likely carrying out these reactions. “IEM” = isoeugenol monooxygenase, “VDH” = vanillin dehydrogenase, “P2M” = phenol 2-monooxygenase, “SH” = salicylate hydroxylase, “4HM” = 4-hydroxybenzoate 1-monoxygenase, “PGR” = phloroglucinol reductase, “PIR” = quercetin 2,3-dioxygenase, “4H1M” = 4-hydroxyphenylacetate 1-monooxygenase.

**Figure 3:**
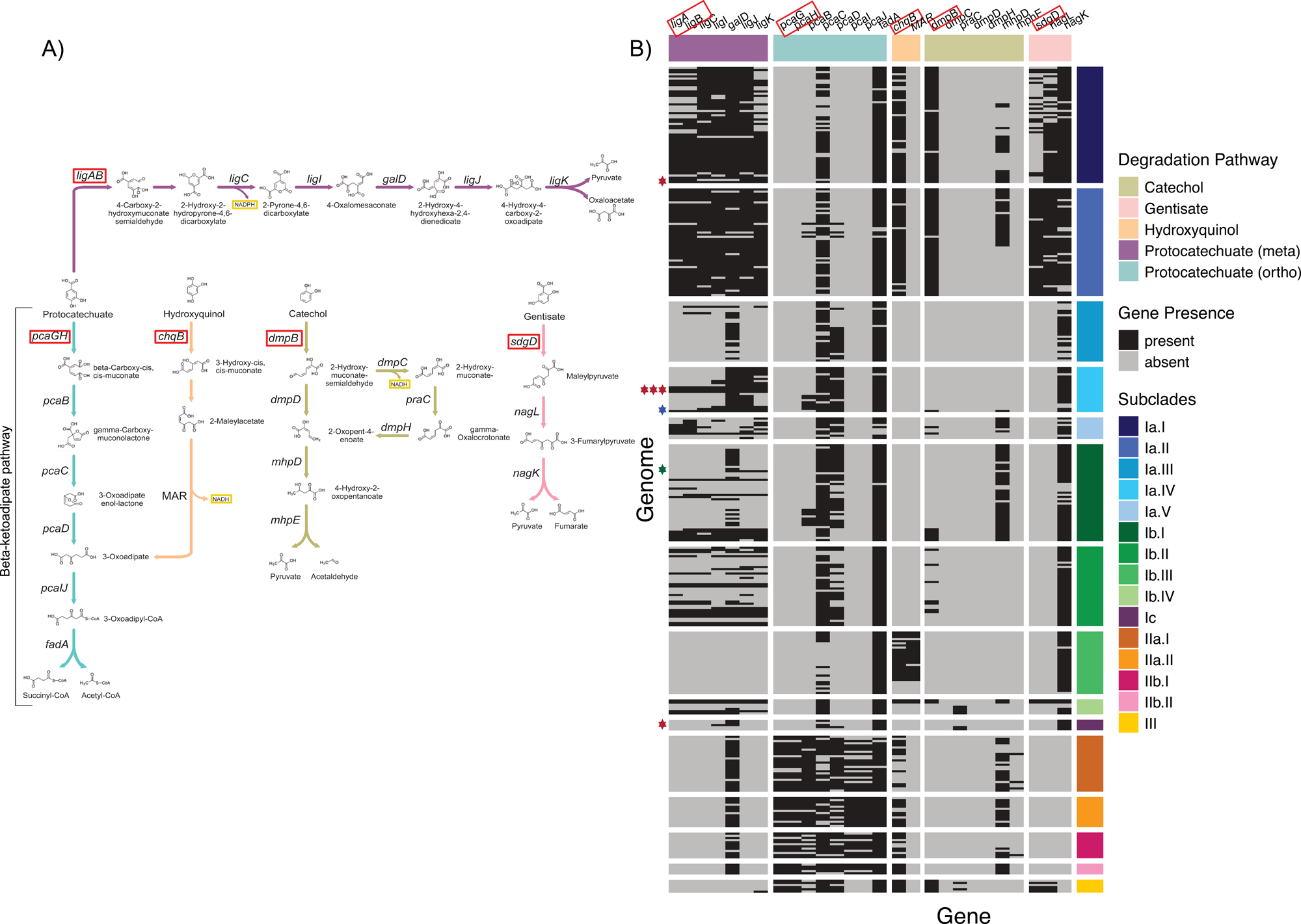
Ring-cleaving dioxygenase genes across the SAR116 clade. (A) Metabolic pathway map. Reaction arrow colors match the metabolic pathway color represented in the heatmap. (B) Presence/absence heatmap of aromatic ring-cleaving dioxygenase genes. Every row is a genome grouped by subclade and every column is a gene grouped by pathway. Stars to the left of the heatmap indicate isolated strains. Red stars indicate LSUCC isolates, the blue star is IMCC1322.

Pyrogallol and phloroglucinol degradation via PgthAB (pyrogallol-phloroglucinol transhydroxylase) and PGR (phloroglucinol reductase), respectively, do not require oxygen (Fig. 1B) [68], but these enzymatic reactions have been detected among aerobic organisms [97], so this metabolism is not restricted by oxygen. Both pyrogallol and phloroglucinol can be produced by marine brown alga species in coastal environments [98]. We found predicted homologs for *pgthAB* across the SAR116 clade, except for subclades Ib.II, Ib.III, and Ib.IV (Fig. 1B), whereas PGR only occurred in subclades Ib.I, Ib.IV, II, and III (Fig. 1B). Dihydrophloroglucinol can be linearized for further metabolic processing via dihydrophloroglucinol cyclohydrolyase [99], however we did not find evidence for this gene in any genomes. Rather than using the carbon from these substrates as a carbon source, these conversions may represent a detoxification strategy (Fig. 2), as dihydrophloroglucinol is less toxic than phloroglucinol [100]. Additionally, these may be examples of metabolic division of labor within the prokaryotic community for polyphenolic degradation, as described in other habitats [101, 102].

Transformations for another polyphenol chemical class, flavonoids, were well represented across subclades Ia.V, IIb, and III (Fig. 1B). We predicted the degradation capacity of flavanones, flavones, and isoflavonoids via the Sam5 flavonoid hydroxylase, a monooxygenase with broad substrate specificity that hydroxylates substrates to weaken the aromatic structure in preparation for ring-cleavage [103]. Sam5 can hydroxylate *p*-coumaric acid, resveratrol, pinostilbene, pterostilbene, naringenin, eriodictyol, apigenin, luteolin, genistein, and daidzein [103–105], all of which marine phytoplankton produce [106–109]. Quercetin is a flavonoid produced by the marine sediment bacterium *Streptomyces fradiae* [54], as well as several microalgae [48, 106] and cyanobacteria [46]. We found homologs for the protein responsible for the degradation of quercetin, quercetin 2,3-dioxygenase, in every SAR116 subclade and encoded in more than 80% of genomes in Ib.III, Ia.IV, and IIb.II (Figs. 1B, 2). To our knowledge, quercetin degradation has not been well-documented in marine systems, although it has strong selective effects in gut microbiomes [110, 111]. The widespread conservation of the quercetin 2,3-dioxygenase implies that quercetin is readily available in marine environments, and likely another key *in situ* metabolite for SAR116. The degradation product, 2-protocatechuoyl-phloroglucinol carboxylic acid, can be further cleaved by a depsidase or esterase [112, 113] into phloroglucinol and protocatechuate (Fig. 2). Since the depsides/esterases for this conversion are not well characterized, we could not confidently predict the cleavage process in SAR116. Nevertheless, our findings suggest marine quercetin degradation deserves further study.

The distributions of genes and pathways related to degradation of phenolic acids had the greatest variation across SAR116 subclades (Fig. 1B). Members of subclades Ia.I, Ia.II, and Ib.IV had the highest representation of phenolic acid degradation pathways. The predicted 4-hydroxybenzoate-CoA ligase (HbaA) for conversion of 4-hydroxybenzoate and benzoate to benzoyl-CoA [114] was widely conserved across subclades Ia.II and IIb (Figs. 1B, 2). The HbaA enzyme does not require oxygen and is commonly described as an anaerobic metabolism [114, 115], however, this enzyme can still function in aerobic systems [115]. Potential for conversion of 4-hydroxyphenylacetate to homoprotocatechuic acid by a 4-hydroxyphenylacetate 3-monooxygenase (HpaB) was encoded across most of the SAR116 clade, with the highest representation in subclades IaV, IIb, and III (Fig. 1B).

Genomes in subclades Ia.I, Ia.II, Ib.IV, and III encoded the pathway to convert ferulic acid to vanillate via feruloyl-CoA synthase but were missing the enzyme for the second step, feruloyl-CoA hydratase/lyase, that produces vanillin (Table S5). However, non-specific enoyl-CoA hydratase/lyases are substitutes for feruloyl-CoA hydratase/lyase [116, 117], and we found many of these genes in SAR116 genomes (Table S2). Although the vanillate conversion to protocatechuate was only occasionally detected via CAMPER (Fig. 1B), manual curation clarified that the capacity for conversion of vanillate to protocatechuate was likely possible across subclade I via vanillate O-demethylase (*vanA*) (Fig. 2, Table S4). Similarly, our manual curation identified a salicylate hydroxylase for salicylate conversion to catechol across subclades Ia, Ib, and II (Fig. 2, Table S4).

The degradation pathways of non-polyphenolic aromatics had the least representation across SAR116 genomes (Fig. 1B), suggesting that polyphenols are more significant metabolites to this group than are other classes of aromatics. That said, we found the transformation pathway of benzoate to protocatechuate in most of the clade, with the most representation in subclades Ia.I, Ia.II, II and III. However, in this two-step pathway, SAR116 genomes lacked homologs of the gene for converting benzoate to 4-hydroxybenzoate, benzoate 4-monooxygenase, but they did have a predicted p-hydroxybenzoate 3-monooxygenase (*pobA*) for converting 4-hydroxybenzoate to protocatechuate (Fig. 2, Table S5). The degradation of benzoyl-CoA, the product of *hbaA*, via the *boxABC* gene suite was primarily found in subclade IIb (Figs. 1B, 2). All SAR116 genomes were missing *boxD*, the gene for 3,4-dehydroadipyl-CoA semialdehyde dehydrogenase, however, it is possible that a promiscuous or non-specific semialdehyde dehydrogenase could be taking the place of *boxD* to finish the pathway. Interspersed throughout SAR116 genomes we found genes for the degradation of small monocyclic aromatic compounds such as m-cresol, and phthalic acid (Figs. 1B, 2). The potential for conversion of m-cresol to 3-methylcatechol via phenol 2-monooxygenase was present in the majority of subclade Ia.I and Ia.II genomes and in some members subclades Ia.IV, Ia.V, and Ib.IV, however, we could not confirm further degradation pathway steps (Figs. 1B, 2).

Members of subclades Ia.II, Ia.IV, and Ic carried *nahAb*, encoding the ferredoxin subunit of napthalane 1,2-dioxygenase (Table S4). Previous work reported an enrichment of SAR116 cells after ^13^C-naphthalene incubations [35]. In those incubations, SAR116 were not considered primary degraders of naphthalene, however they incorporated carbon from naphthalene. Our current genetic evidence suggests that SAR116 cannot degrade naphthalene because they don’t encode the catalytic component of naphthalene 1,2-dioxygenase (Table S4). SAR116 genomes did have genes for the transformation of the downstream substrates of naphthalene degradation such as salicylate and catechol (Table S4, Fig. 3). Therefore, we think it likely that SAR116 metabolize the degradation byproducts of naphthalene metabolism supplied by other taxa. This hypothesis also corroborates observations that SAR116 members are not among the taxa that typically increase in response to oil spills [38, 40, 41, 118, 119]. It is unlikely that SAR116 are degrading polyaromatic hydrocarbons (PAHs) *in situ*, but rather monocyclic aromatics and polyphenols, some of which can result from PAH degradation.

### Ring-cleaving dioxygenases

Polyphenolic and aromatic compounds are funneled through a series of transformations to a subset of substrates that become linearized by a ring-cleaving dioxygenase before further metabolization [120, 121] [122]. There are two families of ring-cleaving dioxygenases: intradiol and extradiol [123]. These two enzyme families can act on the same substrate but yield different products due to the different bonds where cleavage of the aromatic ring occurs, resulting in distinct degradation pathways of the cleaved products (Fig. 3A). Of the polyphenol substrates profiled by CAMPER [42] in SAR116, many of them funnel to protocatechuate degradation [120, 124–129] (Fig. 2) and then to central carbon metabolism (Fig. 3A). A smaller subset of polyphenol substrates ends up as hydroxyquinol, catechol, or gentisate (Fig. 2) [120, 124, 125, 130–133]. Some precursor polyphenols overlap between the cleaved substrates (Fig. 2). For example, 4-hydroxybenzoate degradation can lead to either protocatechuate or gentisate, however, SAR116 members likely convert 4-hydroxybenzoate to protocatechuate via *pobA* (Fig. 2, Table S4). Most SAR116 subclades had predicted ring-cleaving dioxygenases for protocatechuate (*pcaGH, ligAB*), hydroxyquinol (*chqB*), catechol (*dmpB*), and/or gentisate (*sdgD*) (Fig. 3). Subclades Ia.I, Ia.II, II, and III had the greatest representation of ring-cleaving dioxygenase genes (Fig. 3B), indicating that those subclades likely have access to a wider range of *in situ* aromatic compounds to metabolize than the other groups.

Protocatechuate cleavage genes occurred across the entire SAR116 clade with a distinct evolutionary bifurcation: subclade I exclusively encoded the ortho-cleavage pathway via the protocatechuate 4,5-dioxygenase (*ligAB*), whereas subclades II and III exclusively encoded the meta-cleavage pathway via protocatechuate 3,4-dioxygenase (*pcaGH*) (Fig. 3B). The *ligAB* (extradiol dioxygenase) pathway results in pyruvate and oxaloacetate, plus a molecule of NADPH (via *ligC*), whereas the *pcaGH* (intradiol dioxygenase) pathway ultimately produces succinyl-CoA and acetyl-CoA (Fig. 3A). Members of subclades Ia, Ib.I, Ib.II, and Ib.IV, including isolated representatives LSUCC0226 (Ia.IV, closed genome), LSUCC0396 (Ia.IV, closed genome), LSUCC0719 (Ia.I), LSUCC0744 (Ia.IV, closed genome), and IMCC1322 (Ia.IV), had the complete *ligABCIJK/galD* pathway to degrade protocatechuate and utilize the end-products, pyruvate and oxaloacetate, for central carbon metabolism. These genes were most consistently represented in genomes of subclades Ia.I and Ia.II, whereas no subclade Ib.III genomes encoded genes for the degradation of protocatechuate (Fig. 3).

We predicted some members of subclade II could shunt protocatechuate into central carbon metabolism using the *pcaGHBCDIJ/fadA* pathway, also known as the beta-ketoadipate pathway [120]. Subclade III members lacked the *pcaIJ* complex and some were also missing *pcaB* (Fig. 3). However, activity of PcaIJ, a beta-ketoadipate succinyl-CoA transferase, can be replaced by other CoA transferases [134]. Subclade III genomes encoded GctAB, a glutactonate-CoA transferase, within the same operon as *pcaGH* and the downstream post-cleavage degradation genes, *pcaBCD/fadA* (Fig. 4A). Therefore, we propose that subclade III members can also metabolize protocatechuate into central carbon metabolism using the beta-ketoadipate pathway by substituting the *pcaIJ* complex with *gctAB*. The conservation of these pathways across the SAR116 clade suggests that protocatechuate, and its upstream precursors (Figs. 1B, 2), are likely important carbon sources for the group.

**Figure 4:**
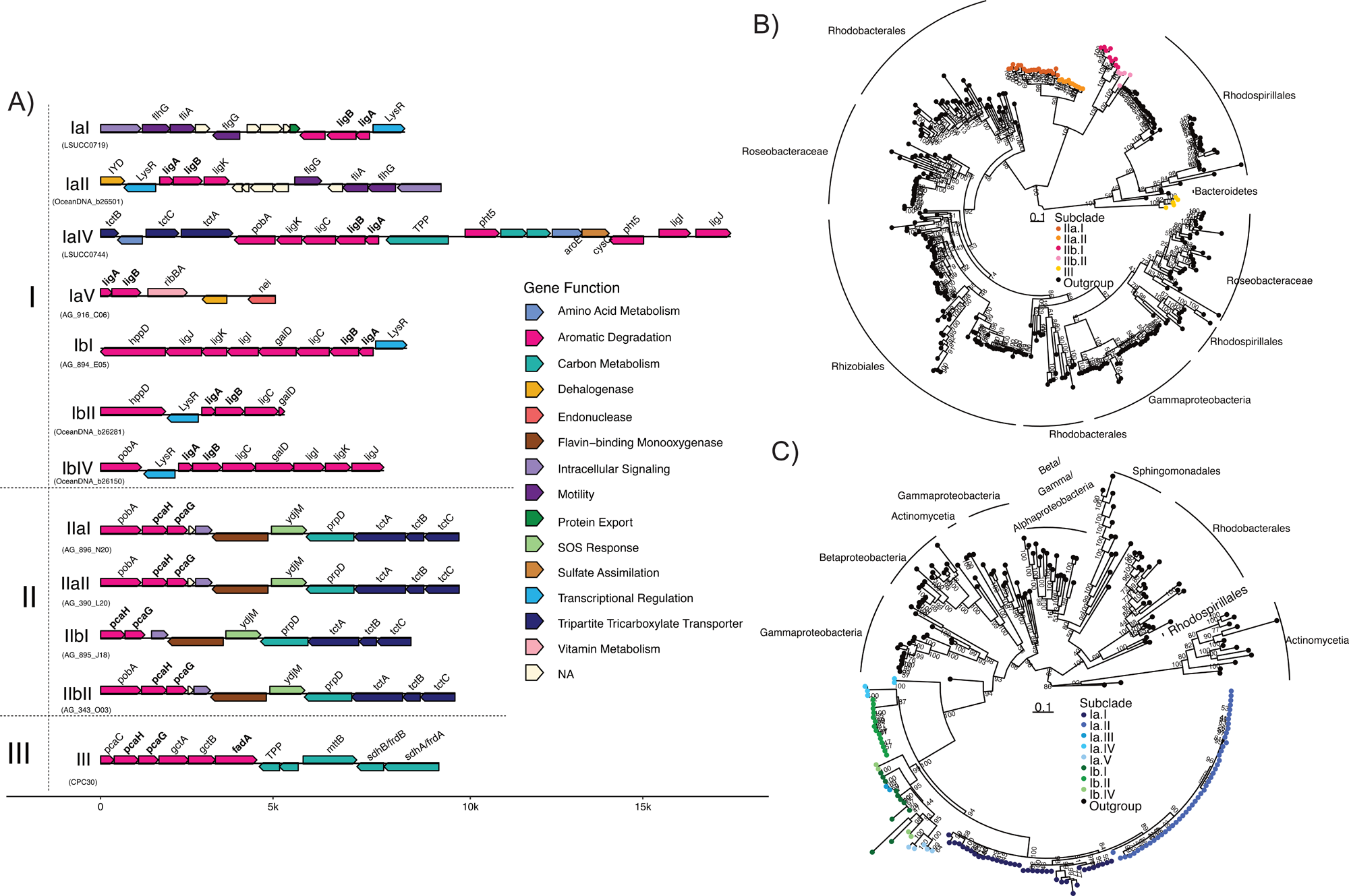
Gene neighborhood and synteny surrounding protocatechuate dioxygenase genes across SAR116 subclades. Each neighborhood is representative of that subclade. (A) Genes are colored by the metabolic function according to the key, with the gene annotation labeled above the gene. Genes that are represented above the contiguous sequence line and pointing to the right are in the forward strand within the genome, and genes below and pointing to the left are in the reverse strand. To the left of the contiguous sequence are the representative genome names and the subclade the genome represents. (B) Phylogenetic tree of PcaH, the protocatechuate 3,4-dioxygenase beta-subunit, with the closest relatives from NCBI RefSeq database represented by black tips. Orange, pink, and yellow tips represent other SAR116 PcaH protein sequences. (C) Phylogenetic tree of LigB, the protocatechuate 4,5-dioxygenase beta-subunit, with closest relatives from NCBI RefSeq database represented by black tips. Blue and green tips indicate other SAR116 LigB protein sequences. Scale bar represents 0.1 changes per position. Bootstrap support values (n=1000) are indicated at nodes.

Gentisate cleavage through *sdgD* was only predicted for subclades Ia.I, Ia.II, Ib.IV, and III, and only a subset of these members in subclades Ia.I, Ia.II, and Ib.IV encoded the complete post-cleavage degradation pathway including *nagL* and *nagK*. Catechol cleavage via catechol 2,3-dioxygenase (*dmpB*), meta-cleavage, was widely distributed across subclades Ia.I, Ia.II, II, and III, with limited representation across other Ia and Ib subclades (Fig. 3B). However, not a single SAR116 genome was predicted to encode a complete pathway for catechol degradation to pyruvate and acetaldehyde (*dmpBD*/*mhpDE*). Thus, the presence of the catechol 2,3-dioxygenase without the post-cleavage degradation pathway may indicate that this enzyme acts in a detoxification role since aromatic compounds are toxic in high concentrations [135, 136]. We predicted the potential for hydroxyquinol cleavage via *chqB* for many genomes in subclades Ia.I, Ia.II, Ib.III, II, and III, but only members of Ib.III and Ib.IV carried a maleylacetate reductase (MAR) required to continue metabolizing the cleaved product and produce NADH (Fig. 3B). However, these groups were not predicted to encode a complete beta-ketoadipate pathway for the remaining steps of degradation (Fig. 3, see below). Therefore, it is likely that hydroxyquinol is not a source of carbon for subclade I members, but rather a potential detoxification mechanism due to the toxicity of hydroxyquinol [137, 138].

### Evolutionary history of ring-cleavage pathways in SAR116

The exclusive presence of meta-cleavage pathway via *ligABCIDJK* in subclade I, vs. the ortho-cleavage pathway via *pcaGHBCDI/fadA* in subclades II and III raised the question as to which of these pathways were ancestrally- or horizontally acquired. Although subclade I did not encode the *pcaGH* protocatechuate 3,4-dioxygenase found in subclades II and III, some of the other downstream genes for the ortho-cleavage/beta-ketoadipate pathway did occur in subclade I (Figs. 1B, 3B). The beta-ketoadipate pathway-specific gene *pcaC* [139] occurred in genomes across all of subclade I, and some genomes also had other beta-ketoadipate specific genes, *pcaB* and *pcaD* [139]. The presence of these genes in subclade I implies that *pcaGH* was lost from this lineage, with substitution of genes for the ortho-cleavage pathway (*ligAB*) occurring prior or subsequently. The phylogenetic tree of PcaH, the active site-containing subunit for the PcaGH complex in subclades II and III, showed a polyphyletic structure where subclade II was sister to members of the *Rhodobacterales* and subclade III were sister taxa to two *Rhodospirillales* clades with four *Bacteroidetes* sequences branching in between (Figs. 4B, S1). Although subclades II and III were not sister clades like they were in the SAR116 species tree [28], the PcaH tree nevertheless suggests an alphaproteobacterial origin and likely vertical inheritance of *pcaGH* because subclades II and III branched near each other, the *Rhodospirillales* were close to SAR116 in larger alphaproteobacterial trees [26, 28, 140], and most of the neighboring groups (e.g., *Rhodobacterales*, *Rhizobiales*) were also in the Alphaproteobacteria. Furthermore, the gene neighborhoods surrounding *pcaGH* showed synteny across subclades II and III (Fig. 4A), which also supported a vertical inheritance hypothesis.

In contrast, the phylogenetic tree for LigB, the active site-containing subunit for LigAB, suggested a different evolutionary history for this pathway. SAR116 subclade I LigB sequences were monophyletic, but the sister taxa were members of the *Gammaproteobacteria* (Fig. 4C), which was incongruent with SAR116’s placement in the *Alphaproteobacteria* (Fig. S2) [28]. Additionally, there was no synteny surrounding the *ligAB* genes in subclade I genomes (Fig. 4A), suggesting that *ligAB* was not vertically inherited, but instead horizontally transferred to the SAR116 clade, and possibly subject to frequent genomic rearrangement thereafter. Additional supporting evidence of this evolutionary history comes from the phylogenies of proteins within the neighborhood of *ligAB.* LysR (Figs. 5A, S3), a transcriptional regulator for benzoate degradation [141], and LigJ (Figs. 5B, S4), a downstream enzyme involved in metabolizing the cleaved protocatechuate product, both also branched sister to the *Gammaproteobacteria*, providing additional support for the hypothesis that the *lig* gene system was horizontally transferred to SAR116 subclade I from *Gammaproteobacteria*.

**Figure 5:**
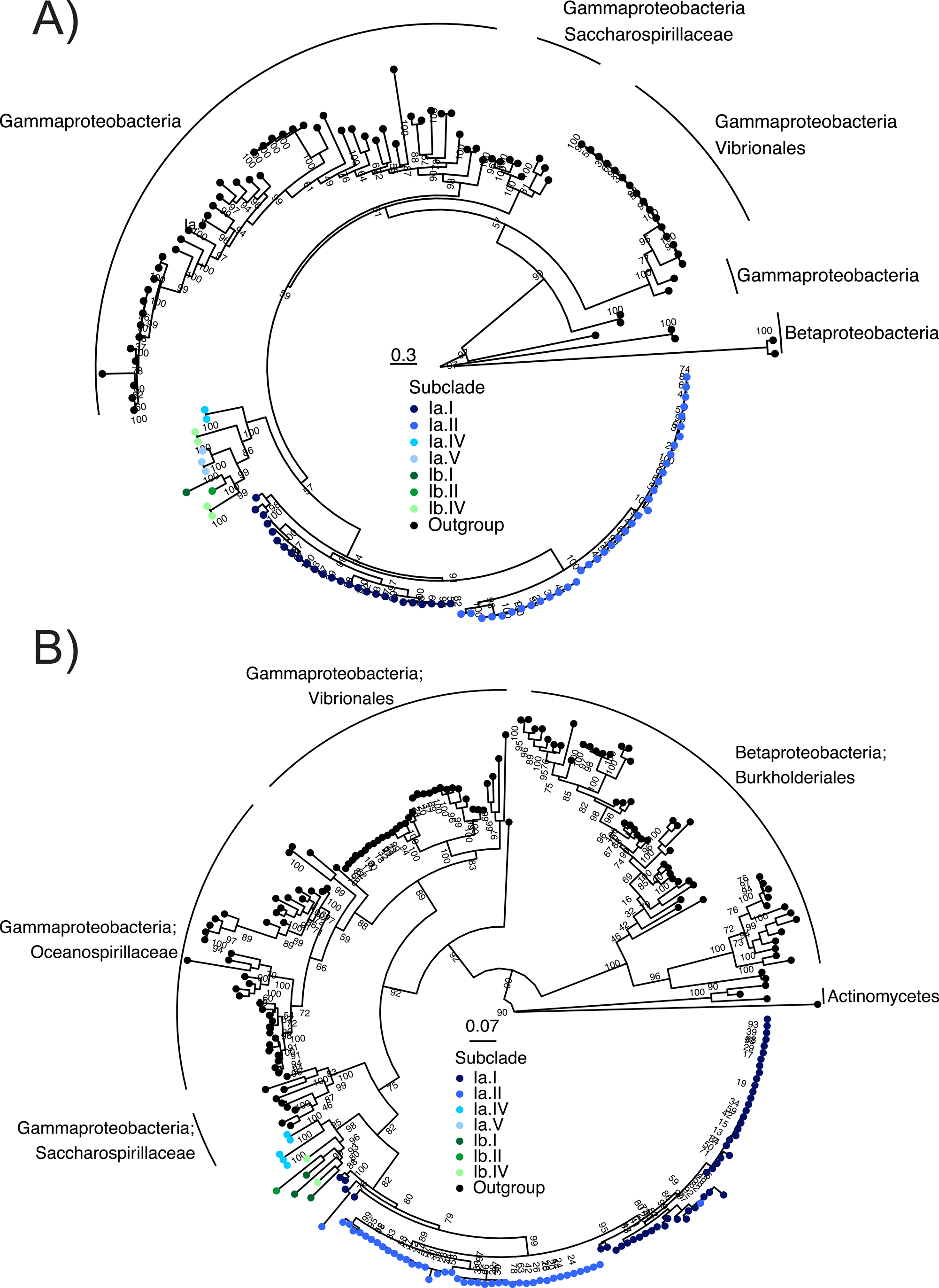
Phylogenetic trees of LysR and LigJ. (A) Phylogenetic tree of transcriptional regulator LysR, and (B) phylogenetic tree of protocatechuate post-cleavage degradation pathway enzyme LigJ. Closest relatives from the NCBI RefSeq database represented by black tips. Blue and green tips indicate SAR116 LysR (A) or LigJ (B) protein sequences. Scale bar represents 0.3 changes per position (A), and 0.07 changes per position (B). Bootstrap support values (n=1000) are indicated at nodes.

Horizontal gene transfer of the meta-cleavage pathway emphasizes the importance of maintaining protocatechuate metabolism for SAR116, since the ancestral Subclade I lineage acquiring this pathway underwent substantial risk with such substantial gene insertions. It is possible that the acquisition of the *ligABCIJK* gene suite occurred prior to the loss of *pcaGH* in Subclade I, thus offering functional redundancy and a possible competitive advantage, after which many of the ortho-cleavage pathway genes could have been lost. Subsequently, many lineages within Subclade I lost some or all of the meta-cleavage pathway. This hypothetical evolutionary path would explain the spotty patterns of both meta- and ortho-cleavage pathway genes in Subclade I members outside of Ia.I and Ia.II. But regardless of the actual chain of events, Subclade I underwent a different evolutionary process in selecting for protocatechuate degradation than Subclades II and III. Further investigation into the kinetics, energetics, and assimilation dynamics of the meta- vs. ortho-cleavage pathways, and the upstream transformations, could help understand whether/how they confer different advantages to their host organisms, and why this metabolism is conserved in so many SAR116 members.

Phylogenies for the other ring-cleaving dioxygenase genes in SAR116 genomes support the hypothesis that all three, *sdgD* (Gentisate 1,2-dioxygenase, Fig. S5), *chqB* (Hydroxyquinol 1,2-dioxygenase, Fig. S6), and *dmpB* (Catechol 2,3-dioxygenase, Fig. S7), were likely vertically inherited in SAR116 based on the neighboring groups being other members of the Alphaproteobacteria (*Rhodobacterales, Rhizobiales, Rhodospirillales*), consistent with SAR116 placement in larger alphaproteobacterial trees [26, 28, 140].

### Polyphenol metabolism in host-associated subclade

Evidence suggests that one subclade of SAR116 (Ic) has a host-associated component of its life cycle, as it has been detected in association with corals and sponges and undergoes extensive pleiotropy [28]. Subclade Ic genomes have genes for polyphenol metabolism (Fig. 1), including ring-hydroxylating dioxygenases (Table S4), but without ring-cleaving dioxygenases (Fig. 3B, Table S3). This likely reflects the loss of *pcaGH* observed throughout subclade I but without the corresponding substitution of *ligABCIJK*. Many flavonoids, phenolic acids, tannins, and terpenoids have been detected in coral and sponge extracts [43, 50, 52] they are also produced by the dynamic microbiome that these organisms host [51]. Subclade Ic polyphenol metabolism, even without ultimate ring-cleavage, may benefit corals, sponges, and other microbiome members as a potential detoxification strategy, since the accumulation of polyphenols and phenolic acids can have toxic effects for both the host and microbiome [142–145]. In fact, symbionts carrying phenoloxidases, which are encoded by subclade Ic and many other SAR116 subclades, can aid in coral and sponge innate immunity due to free radical scavenging [146, 147].

### In situ activity

To investigate whether SAR116 members might be using these metabolisms *in situ*, we examined gene expression of ring-cleaving dioxygeneases using metatranscriptomic read recruitment to SAR116 genomes. We focused specifically on the ring-cleaving dioxygenase genes (Fig. 3) because they serve as bottlenecks for carbon acquisition in polyphenolic and aromatic degradation [125] (Figs. 2, 3). We found evidence for expression of all four ring-cleaving dioxygenase genes across coastal and open ocean environments (Fig. 6), suggesting that polyphenolic metabolism occurs widely in SAR116 bacteria. SAR11 protocatechuate-cleaving dioxygenases were expressed across the global surface oceans, with hotspots of activity occurring in specific regions, such as the Mediterranean and Arabian Seas (Fig. 6A), as well as in the deep chlorophyll maximum (DCM), although at lower levels (Fig. 6B). Expression of hydroxyquinol, catechol, and gentisate ring-cleaving dioxygenases also occurred in surface waters and the DCM (Fig 6C-H). We observed relatively little expression of catechol ring-cleaving dioxygenases at the DCM (Fig. 6E). However, hydroxyquinol and gentisate showed pockets of high activity at certain locations within the DCM (Figs. 6D, H). Regardless of the pathway, we found evidence for expression from all major subclades in which it occurred (I, II, and III). These patterns suggest SAR116 bacteria have widespread ongoing interactions with polyphenolic compounds globally that deserve further investigation.

**Figure 6:**
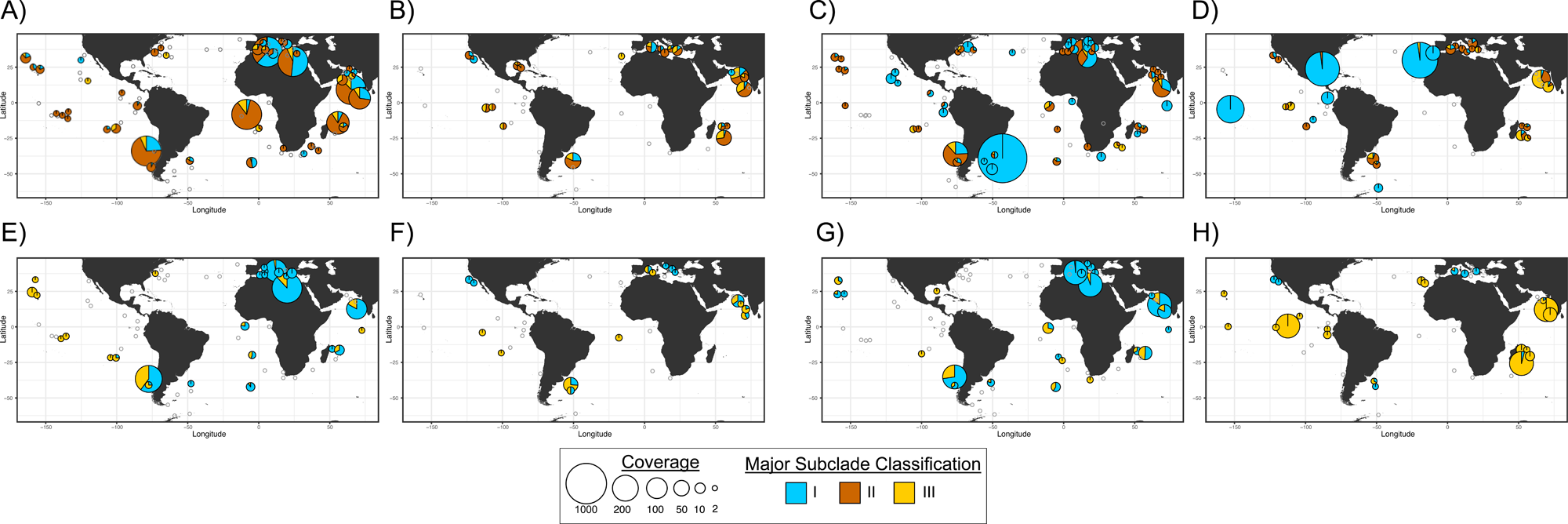
Ecological distribution of ring-cleaving dioxygenase gene expression across SAR116 major subclade classifications. World maps showing the spatial distributions of metatranscriptomic recruitment of protocatechuate ring-cleaving dioxygenase genes at the surface (A), and deep chlorophyll maximum (B). Hydroxyquinol ring-cleaving dioxygenase genes at the surface (C), and deep chlorophyll maximum (D). Catechol ring-cleaving dioxygenase genes at the surface (E) and deep chlorophyll maximum (F). Gentisate ring-cleaving dioxygenase genes at the surface (G), and deep chlorophyll maximum (H). The size of the pie chart represents the total coverage, or number of reads, that mapped to the ring-cleaving dioxygenases. The colors in the pie charts represent the major SAR116 subclade classifications, and the colored proportion of the pie chart represents the proportion of total reads mapped per major subclade classification. The grey unfilled circles represent the metatranscriptomic samples where expression of protocatechuate ring-cleaving dioxygenases were not detected.

Previous discussion of polyphenolic compounds, defined as “humic” or “terrigenous” DOM, have predominantly focused on coastal or estuarine settings [129, 148–150]. While our evidence supports SAR116 using polyphenol metabolism in these ecosystems, we also observed expression of all four ring-cleaving dioxygenases across open ocean sites (Fig. 6). This corroborates observations from open ocean sampling sites in the Pacific and Atlantic oceans, where polyphenols and lignin derivatives constituted 0.7-2.4% [150] and up to 25% [151] of the total surface ocean DOM. Although riverine and terrestrial input accounts for some of this [150, 151], there is also evidence that marine microbes synthesize these compounds *in situ* [43–47, 53]. Our results indicate that SAR116 members likely transform and consume polyphenol compounds regardless of marine ecosystem and coastal proximity (Fig. 6).

## Conclusions

These results provide evidence for extensive polyphenol degradation capacity in SAR116 bacterioplankton, another metabolic niche than what has been previously highlighted [26, 28, 29, 152], and emphasize polyphenols as important compounds in marine systems beyond coastal and estuarine environments. The global distribution of dioxygenase expression patterns, in conjunction with their evolutionary history within the SAR116 clade, strongly suggests that these metabolic pathways are ancient and central processes for these organisms in myriad ocean contexts, rather than just opportunistic responses to allochthonous inputs. In fact, similar metabolic potential in other marine taxa [36, 129, 134, 148, 153–157] suggests that marine microbial communities may be employing a metabolic division of labor for degrading polyphenols, as seen in other systems [42, 101, 102]. The association of SAR116 with phytoplankton blooms [32–34], suggests that polyphenol degradation may connect SAR116 with phytoplankton physiology

Future work investigating polyphenol metabolism in SAR116 and other marine taxa will help define the extent of these processes in marine systems and identify additional important metabolites within the DOM pool. Luckily, SAR116 cultures are available to help validate these bioinformatic hypotheses and evaluate the physiological variation in SAR116 subclades to different polyphenol compounds, including idiosyncrasies in growth rate, biomass, and energy yields [21, 26–28].

## Data availability

Genome assemblies for isolate representatives LSUCC0226, LSUCC0396, LSUCC0684, LSUCC0719, and LSUCC0744 are available on NCBI under CP166132, CP166131, CP166130, JBFPJN000000000, and CP166129, respectively. Raw reads are available under BioProject PRJNA1133775. Supplementary files, including the pangenome summary (Table S2), and R scripts are available on FigShare https://doi.org/10.6084/m9.figshare.31306816.v3. Cultures of the LSUCC isolate representatives used in this analysis are available upon request.

## Acknowledgements

We would like to thank Kelly Wrighton for time and resources in collaboration with the CAMPER analysis. We would also like to acknowledge the Center for Advanced Research Computing at the University of Southern California (https://carc.usc.edu) for computational resources that have contributed to the results in this publication. This work was supported by This work was supported by a Simons Early Career Investigator in Marine Microbial Ecology and Evolution Award, and NSF Biological Oceanography Program OCE-1945279 and Emerging Frontiers Program EF-2125191 grants to J.C.T.

## Supplementary figures

**Figure S1:**
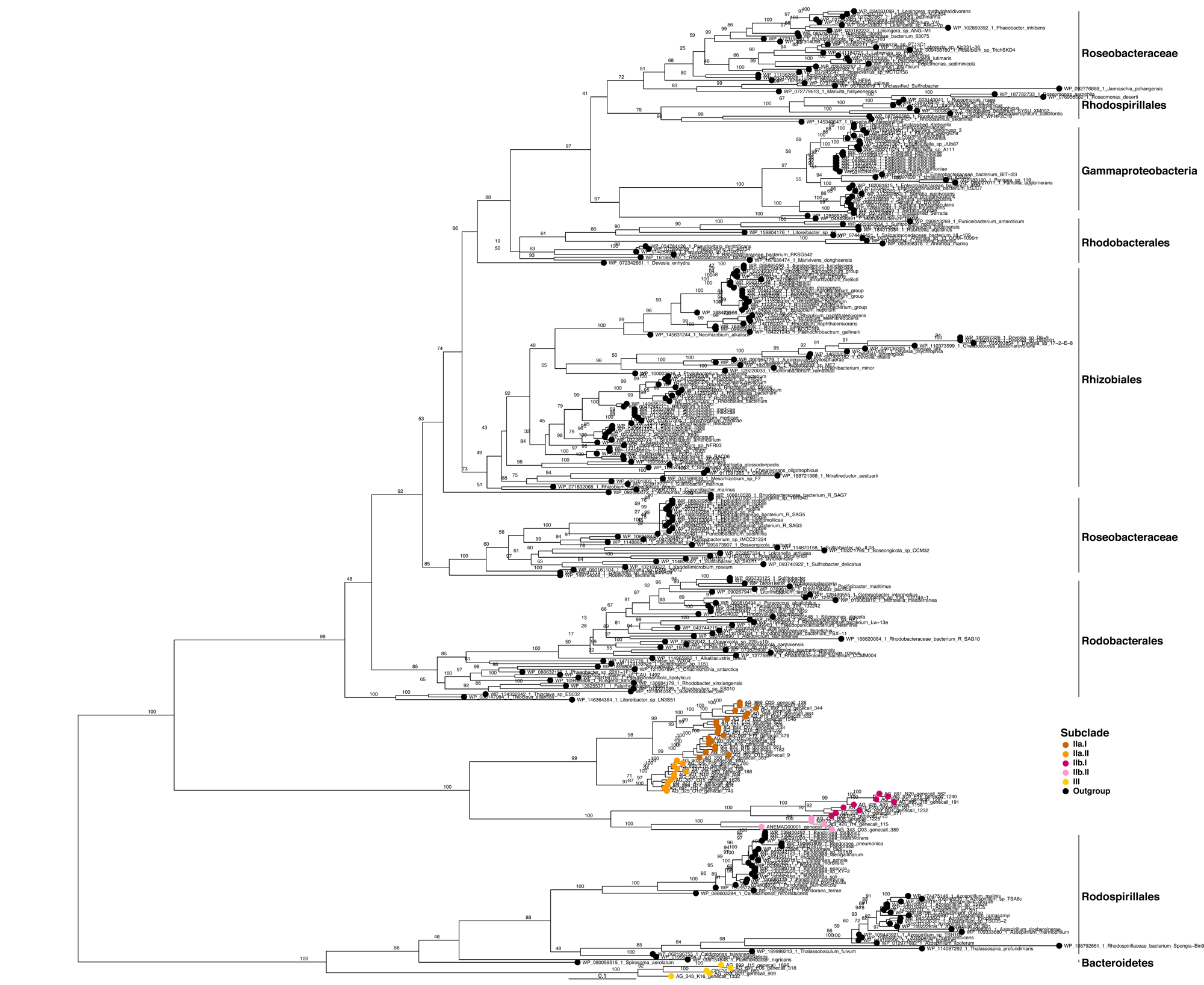
Phylogenetic tree of PcaH. Protocatechuate 3,4-dioxygenase beta-subunit, with closest relatives from NCBI RefSeq database, represented by black tips. Orange, pink, and yellow tips indicate SAR116 PcaH protein sequences. Scale bar represents 0.1 changes per position. The subclade designation can be mapped to the branching structure in the tree using the key. Bootstrap support values (n=1000) are indicated at nodes.

**Figure S2:**
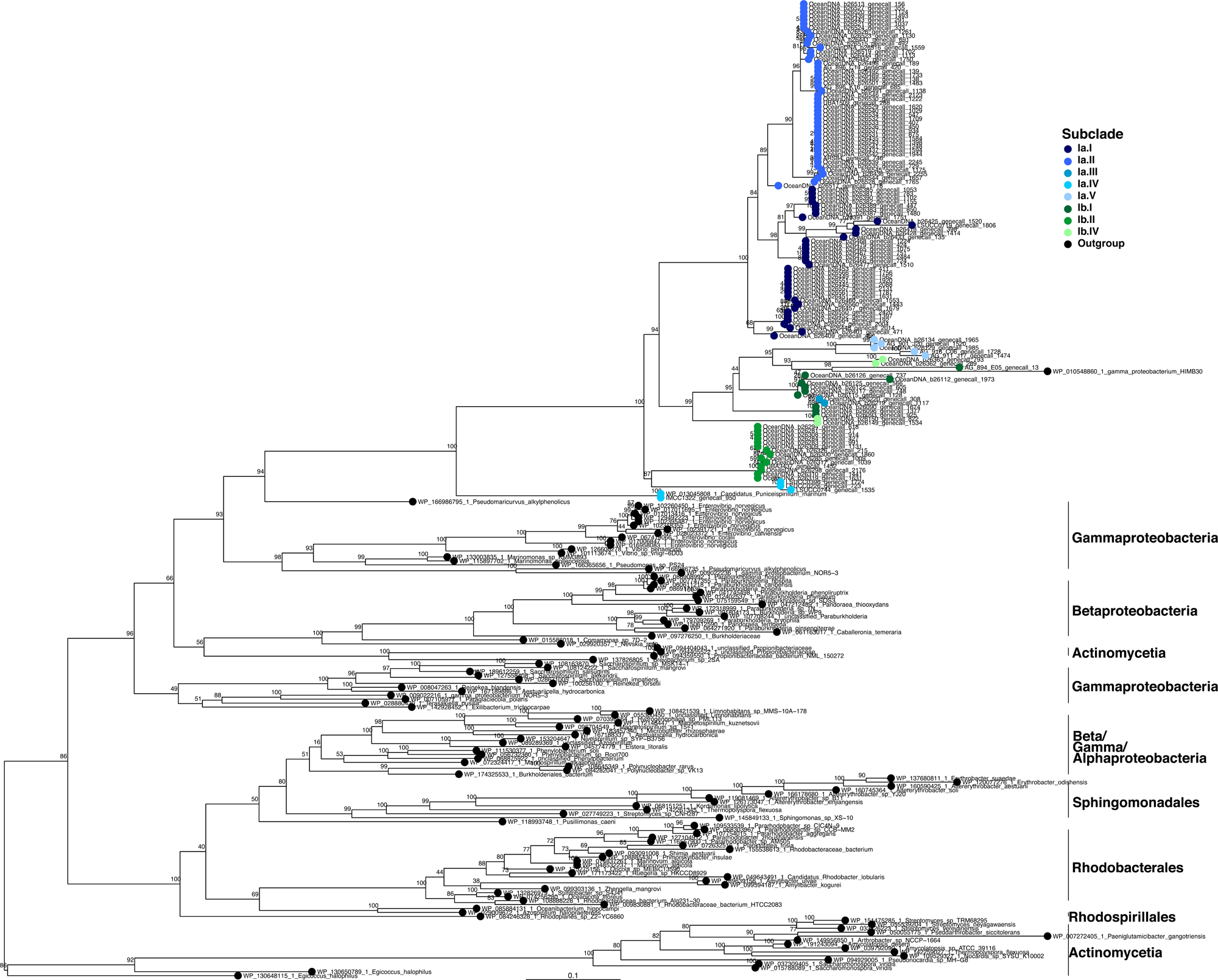
Phylogenetic tree of LigB. Protocatechuate 4,5-dioxygenase beta-subunit, with closest relatives from NCBI RefSeq database represented by black tips. Blue and green tips indicate SAR116 LigB protein sequences. Scale bar represents 0.1 changes per position. The subclade designation can be mapped to the branching structure in the tree using the key. Bootstrap support values (n=1000) are indicated at nodes.

**Figure S3:**
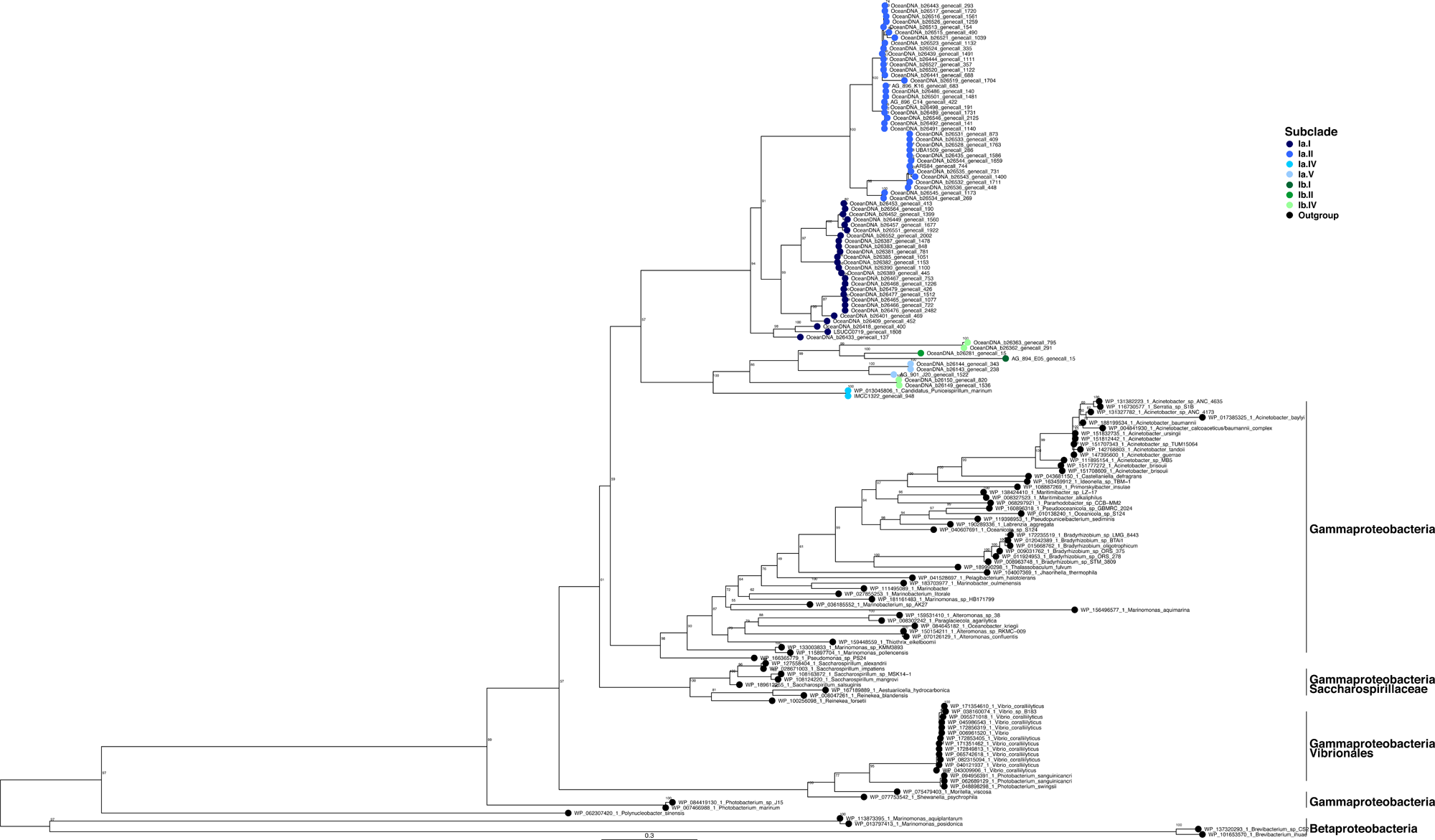
Phylogenetic tree of LysR. Transcriptional regulator LysR, with closest relatives from NCBI RefSeq database represented by black tips. Blue and green tips indicate SAR116 LysR protein sequences. Scale bar represents 0.3 changes per position. The subclade designation can be mapped to the branching structure in the tree using the key. Bootstrap support values (n=1000) are indicated at nodes.

**Figure S4:**
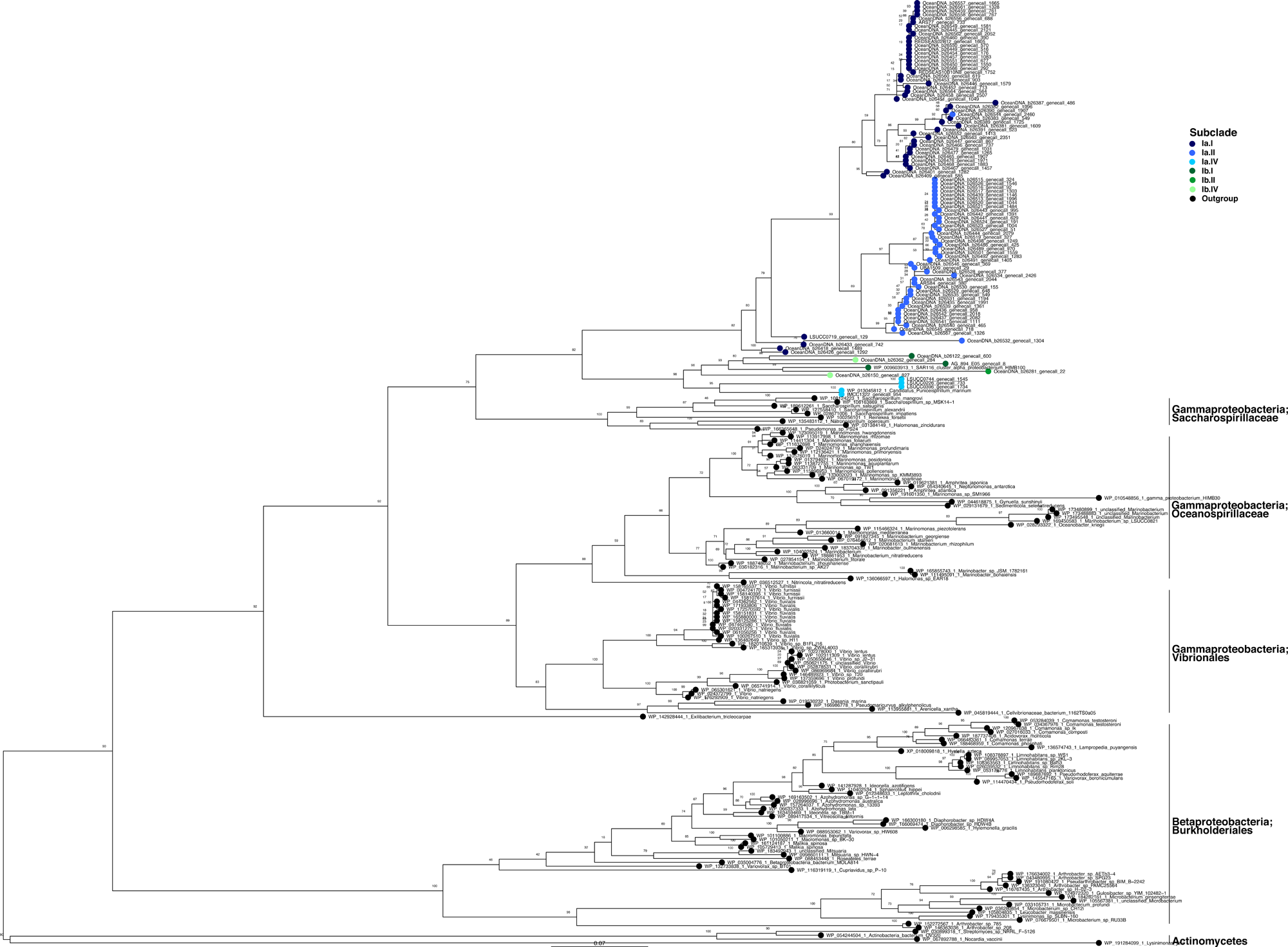
Phylogenetic tree of LigJ. protocatechuate post-cleavage degradation pathway enzyme LigJ, with closest relatives from NCBI RefSeq database represented by black tips. Blue and green tips indicate SAR116 LigJ protein sequences. Scale bar represents 0.07 changes per position. The subclade designation can be mapped to the branching structure in the tree using the key. Bootstrap support values (n=1000) are indicated at nodes.

**Figure S5:**
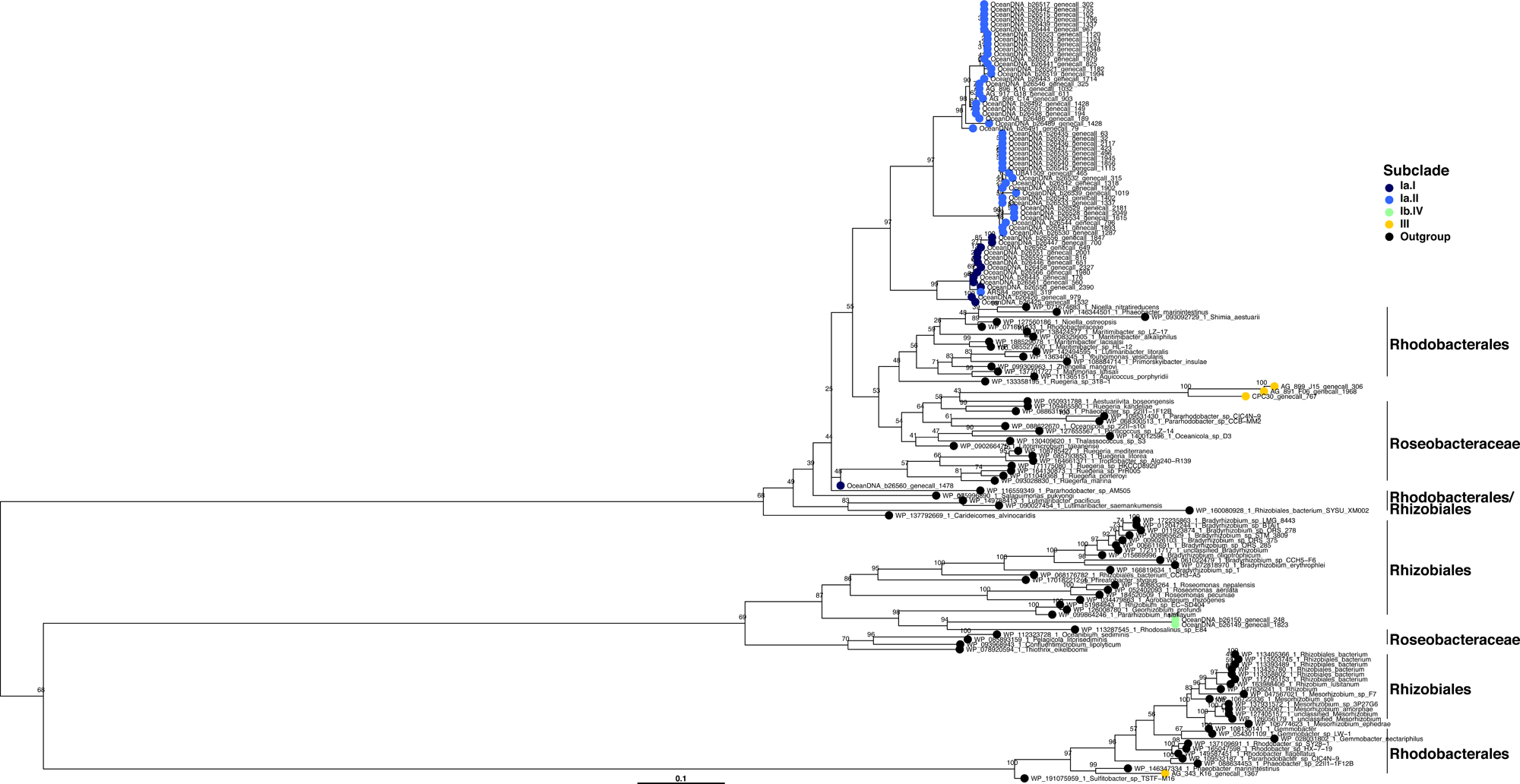
Phylogenetic tree of SdgD. Gentisate 1,2-dioxygeanse, with closest relatives from NCBI RefSeq database represented by black tips. Blue, green, and yellow tips indicate SAR116 SdgD protein sequences. Scale bar represents 0.1 changes per position. The subclade designation can be mapped to the branching structure in the tree using the key. Bootstrap support values (n=1000) are indicated at nodes.

**Figure S6:**
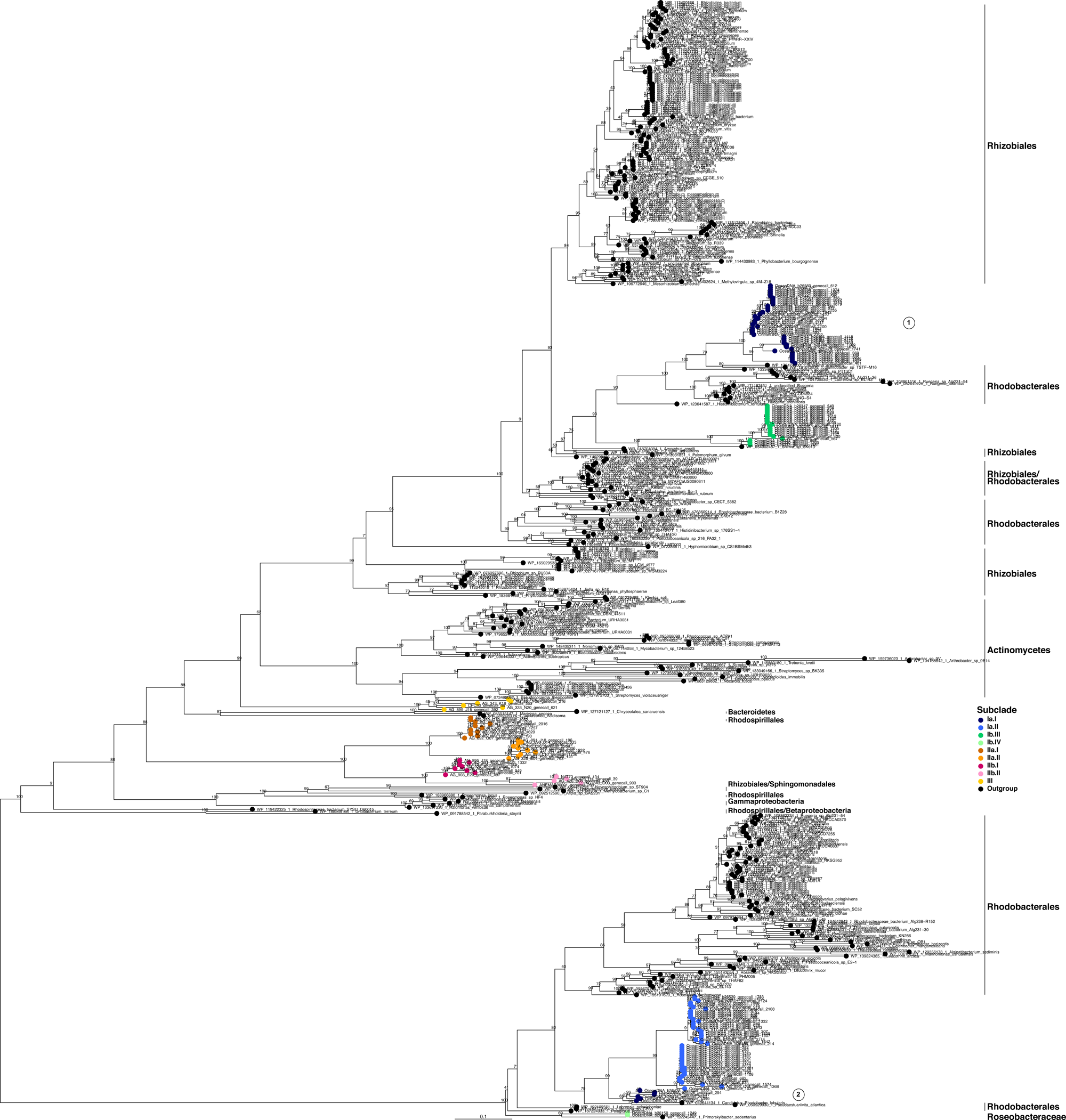
Phylogenetic tree of ChqB. Hydroxyquinol 1,2-dioxygenase, with closest relatives from NCBI RefSeq database represented by black tips. Blue, green, orange, pink, and yellow tips indicate SAR116 ChqB protein sequences. Scale bar represents 0.1 changes per position. The subclade designation can be mapped to the branching structure in the tree using the key. Bootstrap support values (n=1000) are indicated at nodes.

**Figure S7:**
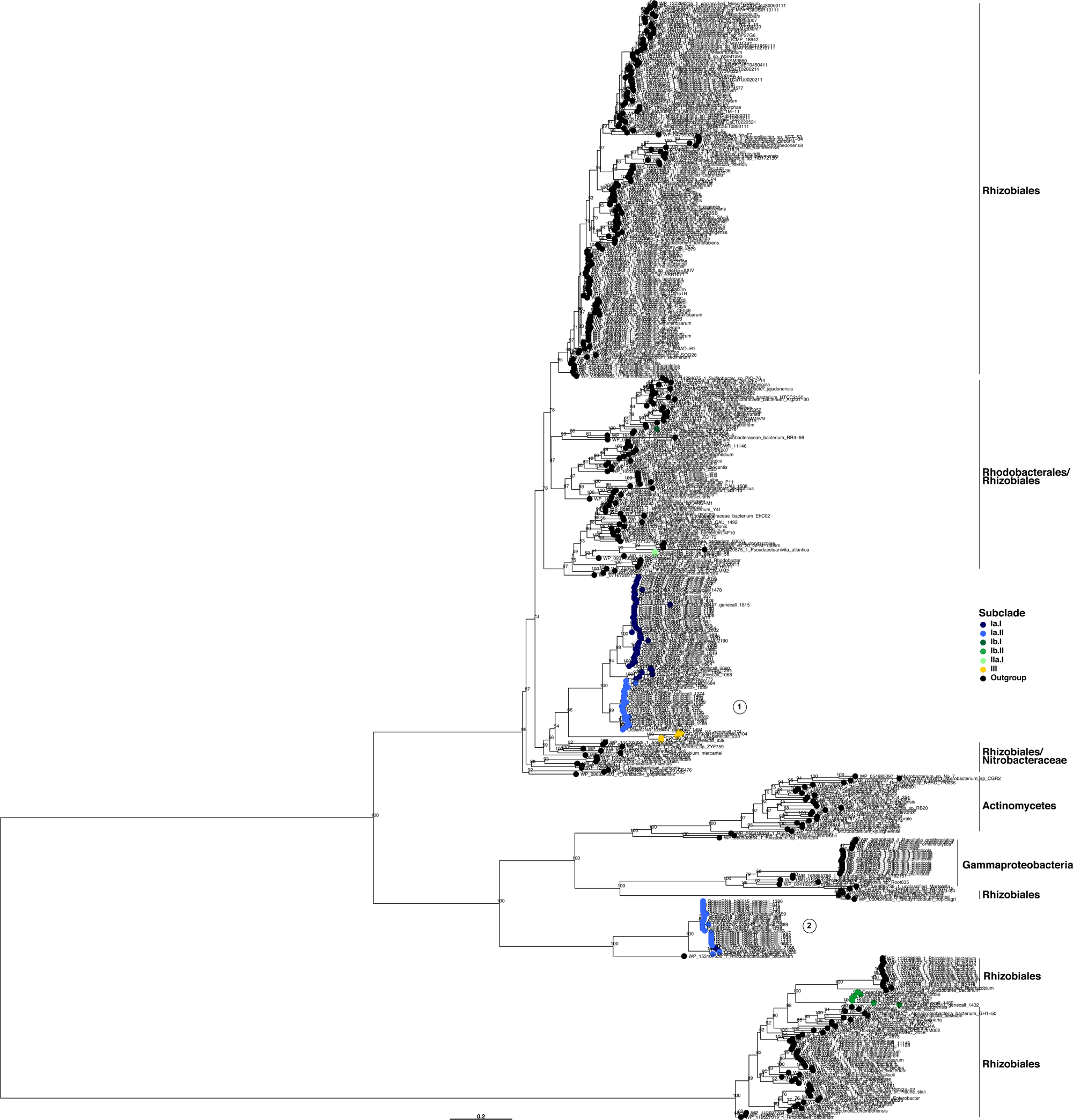
Phylogenetic tree of DmpB. Catechol 2,3-dioxygenase, with closest relatives from NCBI RefSeq database represented by black tips. Blue, green, and yellow tips indicate SAR116 DmpB protein sequences. Scale bar represents 0.2 changes per position. The subclade designation can be mapped to the branching structure in the tree using the key. Bootstrap support values (n=1000) are indicated at nodes.

## Supplementary tables

**Note: due to file size, Tables S2 and S5 are hosted independently at FigShare:** https://doi.org/10.6084/m9.figshare.31306816.v3

**Table S1: Genome information.** Includes accession numbers for SAR116 and outgroup genomes. Also includes CheckM output for all SAR116 genomes, and genome completion summary by subclade.

**Table S2: Complete pangenome summary** of all SAR116 genomes from Anvi’o-7.1. Includes all gene annotations for all genomes, and clustered orthologous groups.

**Table S3: SAR116 ring-cleaving dioxygenase genes.** Subset of annotation from Table S2 that includes ring-cleaving dioxygenase genes in SAR116 genomes.

**Table S4: SAR116 ring-hydroxylating dioxygenase genes.** Subset of annotation from Table S2 that includes ring-hydroxylating dioxygenase genes in SAR116 genomes.

**Table S5: Raw CAMPER annotations.** Direct annotation output from CAMPER.

**Table S6: CAMPER annotation summary table.** Raw annotations were distilled in DRAM to generate the summary table according to degradation pathway modules.

**Table S7: Gene synteny surrounding protocatechuate ring-cleaving dioxygenases.** Annotations on contiguous sequences to identify gene neighborhoods surrounding the protocatechuate ring-cleaving dioxygenases. Annotations subsetted from Table S2 and ordered per genome by their gene-caller-id.

**Table S8: Counts per metatranscriptomic sample per ring-cleaving dioxygenase across SAR116 genomes.** Counts per loci, per genome across all metatranscriptomic samples. Each sheet corresponds to a different polyphenolic substrate for which the SAR116 genomes encode ring-cleaving dioxygenases.

